# Morphological cell profiling for drug repurposing against SARS-CoV-2 infection

**DOI:** 10.1101/2025.08.28.672794

**Authors:** Elin Asp, Jonne Rietdijk, Marianna Tampere, Hanna Axelsson, Duncan Njenda, Swapnil Potdar, Adelinn Kalman, Polina Georgieva, Maris Lapins, Flavio Ballante, Alicia Soler, Martin de Kort, Tero Aittokallio, Andrea Zaliani, Maria Kuzikov, Philip Gribbon, Donald Lo, Jordi Carreras-Puigvert, Brinton Seashore-Ludlow, Ola Spjuth, Päivi Östling

## Abstract

Antiviral drug discovery has traditionally focused on targeting viral proteins, while host-directed strategies remain largely underexplored. Here, present a systematic drug repurposing strategy leveraging morphological profiling to identify host-targeting antivirals. Our image-based approach combines viral protein immunostaining with high-content Cell Painting analysis to simultaneously assess viral replication and provide in-depth analysis of host cell responses. By screening 5,275 repurposable drugs against SARS-CoV-2 infected cells, we identified compounds that reversed the infected cell phenotype, including ones not detected by conventional cytopathicity and antibody-based assays. A counter-screen excluded compounds whose antiviral activity was likely driven by drug-induced phospholipidosis (DIPL). Pathway enrichment analysis of compounds validated by both Cell Painting dose-response and DIPL assays, revealed host processes frequently hijacked by viruses, including innate immune responses and kinases. Among the top hits, both novel candidates, such as serdemetan, and previously reported broad-spectrum antivirals, such as sunitinib, were identified. Our approach constitutes an adaptable and scalable platform suited for diverse viral pathogens and cell systems. We provide a resource of open access screening data, images, and automated analysis pipelines to advance both antiviral discovery and pandemic preparedness.

## Introduction

Over the past centuries, the emergence of pathogenic viruses such as Ebola virus, Zika virus, chikungunya virus, and severe acute respiratory syndrome coronavirus 2 (SARS-CoV-2), has significantly impacted human health, resulting in hundreds of thousands of deaths annually (Baker et al., 2022; Cenciarelli et al., 2015; Mayer et al., 2017). Several factors, including increased human and domestic growth, climate change, and global trade, have led to an increase in the number of viral outbreaks (Bogich et al., 2012; Carlson et al., 2022; Smith et al., 2009). Mutation-prone viruses–especially those with RNA genomes–evolve rapidly, increasing the risk of acquiring traits that enhance human infection or zoonotic spillover (Keusch et al., 2022). In the event of a future pandemic, rapid methods for identifying effective antiviral agents against novel or re-emerging viral variants are vital.

Antiviral drugs represent an important means of slowing the spread of viruses during new outbreaks, as they can be rapidly deployed, are effective even after infection, and are suitable for immunocompromised individuals who cannot be vaccinated (Edwards et al., 2022; Neumann and Kawaoka, 2023). Small-molecule antivirals have already proven their value by reducing morbidity and mortality in infectious diseases such as hepatitis B, hepatitis C, HIV, and herpes simplex (De Clercq and Li, 2016). There are two main classes of antiviral agents: direct-acting antivirals (DAAs) and host-directed antivirals (HDAs). While DAAs are most widely used, they are limited by the rapid emergence of resistance and typically lack broad-spectrum activity. In contrast, HDAs, which target host proteins involved in the virus life cycle, tend to offer broader spectrum potential and a higher barrier to resistance, making them effective against a wide variety of viruses. These advantages, however, come at the cost of a higher risk of off-target effects from treatment (Geraghty et al., 2021; Ji and Li, 2020; Kumar et al., 2020).

The most common assay format for phenotypic antiviral drug screening is based on the indirect measurement of viral infection through virus-induced cytopathic effect (CPE). In this approach, compounds are assessed for their ability to prevent cell death caused by viral replication, serving as a proxy for antiviral activity. While widely used due to its simplicity and scalability, this method captures primarily end-stage infection outcomes, missing early and subtle antiviral effects, and is limited to lytic viruses (Bernatchez et al., 2018). In contrast, antibody-based screens, using virus-specific antibodies to track viral infectivity, can directly quantify viral proteins or particles, enabling detection of infection earlier in the viral life cycle. However, these assays typically neglect the effects of the compounds on the host cell.

To better capture host-cell responses, morphological profiling is emerging as a powerful phenotypic screening approach for antiviral research (Doijen et al., 2024; Goethals et al., 2025, 2025; Heiser et al., 2020). Morphological profiling is a high-throughput, image-based approach that extracts thousands of features from single cells, to create high-dimensional morphological profiles. The most widely adopted method is the Cell Painting assay, which involves exposing cells to perturbations, staining key organelles with multiplexed fluorescent dyes, and imaging across multiple fluorescent channels (Bray et al., 2016). This approach captures rich, multivariate data in a single, cost-effective assay and has recently gained significant traction in the field of drug discovery (Seal et al., 2025). To this end, we previously adapted the Cell

Painting assay for antiviral drug screening by combining morphological profiling with antibody-based detection of virus-infected cells (Rietdijk et al., 2021). This enables tracking of viral infection, identifying infection-specific morphological signatures and screening for compounds that reverse virus-induced phenotypes, offering insights into both antiviral activity, host-cell health and putative mechanisms of action of the compounds.

Drug repurposing has emerged as a popular screening strategy, with the potential to shorten the path for bringing a drug to the market due to the known pharmacological and human safety profiles (Ashburn and Thor, 2004). Several drug repurposing efforts were conducted during the COVID-19 pandemic, identifying numerous promising agents against SARS-CoV-2 (Chen et al., 2021; Ellinger et al., 2021; Mirabelli et al., 2021; Riva et al., 2020). However, the urgent nature of the pandemic overlooked confounding factors, such as drug-induced phospholipidosis (DIPL). Recent studies have suggested that the antiviral activity of several of these drugs was predominantly driven by drug-induced phospholipidosis (DIPL), rather than by specific engagement of viral or host targets (Tummino et al., 2021). This has led to the identification of compounds with non-specific *in vitro* antiviral activity rather than target-based activity with relevance *in vivo* (Tummino et al., 2021). As such, assessing DIPL is an important measure in antiviral drug screening to ensure target-based activities.

Here, we present a strategy for systematic drug repurposing based on morphological profiling to identify host-targeting antiviral candidates against SARS-CoV-2. Using a drug repurposing library of 5,275 compounds, we assessed their ability to reverse host cell morphology toward the uninfected state and compared the results with traditional antiviral read-outs: protection against virus-induced CPE and antibody-based detection of viral replication. We then validated hits from the three readouts in dose-response and deprioritized DIPL-inducing compounds, revealing several promising host-directed antiviral candidates with low DIPL. To further guide compound prioritization, we integrated a target product profile (TPP), which outlines the desired attributes of a compound for clinical use (“Target product profiles,” n.d.). We provide here open access to a valuable collection of high-quality screening datasets generated across multiple assay platforms, including viability measurements across five human cell lines, a CPE screen, DIPL across several concentrations and timepoints, and high-content data from morphological cell profiling and antibody-based detection. We believe this work provides a valuable framework and resource to accelerate the discovery of future host-directed antivirals.

## Materials and methods

### Biosafety

Experiments involving infectious viruses were performed under biosafety laboratory conditions according to Swedish Work and Health Authorities. Work with SARS-CoV-2 was conducted under BSL-3 conditions and work involving HCoV-229E and Zika virus was conducted under BSL-2 conditions.

### Cell lines

African green monkey kidney cell line Vero E6 was maintained in Dulbecco’s modified eagle medium (DMEM) high glucose (Gibco) and supplemented with 1% L-glutamine (Cytiva). Validation experiments were performed using the human lung alveolar cell line A549^ACE2^, which was genetically transduced to constitutively overexpress the ACE2 receptor, developed and donated by the Oscar Fernandez-Capetillo lab (Porebski et al., 2024). The cells were maintained in DMEM/F-12 (Fisher Scientific) with GlutaMAX supplement and 1x non-essential amino acids (Cytiva). Cell lines used in the viability screen included Calu-3 and Caco-2, maintained in MEM with Earl’s salts (Sigma) supplemented with non-essential amino acids (Cytiva); U2OS, cultured in McCoy’s 5A medium (Sigma); and HepG2, maintained in high-glucose DMEM. All cell lines were additionally supplemented with 10% fetal bovine serum (Gibco) and 1% penicillin/streptomycin (Cytiva). Additional cell culturing details are described in supplementary methods.

All cells were kept at 37 °C in a humidified atmosphere with 5% CO_2_. They were detached using TrypLE Express (Gibco), counted, and diluted to appropriate densities with their respective media. Prior to experiments, cells were tested for mycoplasma using a luminescence-based assay (Lonza), and the cell lines were verified using STR profiling.

### Compound library and compound spotting

Compound handling was performed by the SciLifeLab Compound Center (CBCS, Solna, Sweden). The drug repurposing library was sourced from SPECS in DMSO at a concentration of 10 mM. Mechanism-of-action annotations for the full library were obtained from the Clue.io Repurposing Hub unless otherwise noted (Corsello et al., 2017). In brief, chemicals were dispersed between 2.5 nl and 40 nl using the Echo 55/6500 (Labcyte) liquid handler into 384-well assay plates (Phenoplate, Revvity) and stored at -20 °C prior to experimentation. For the full repurposing library (n = 5275), compounds were distributed over the plates with one technical replicate and two biological replicates. The validation screen by Cell Painting in A549^ACE2^ was performed in a 4 point-dose-response in concentrations ranging from 0.1 to 30 μM, in three replicates placed on separate multiwell plates. To reduce bias by positional effects in the microwell plates, the conditions were distributed over the plates using Plate Layouts using Artificial Intelligence Design (PLAID) (Francisco Rodríguez et al., 2023).

### Virus infection

The SARS-CoV-2 strain (CoV-2/human/SWE/01/2020) was obtained from the Public Health Agency of Sweden and propagated in the Vero E6 cells. Viral titers were determined by plaque assay and prepared stocks were stored at -80 °C. Cells were infected in suspension containing serum-free media at multiplicity of infection (MOI) of 0.05 for one hour at 37 °C and 5% CO_2_ atmosphere. Following infection, the cells were centrifuged at 1000 rpm for 5 min, washed with PBS, centrifuged, and resuspended in complete cell culture media. The cells were seeded at an appropriate cell density in pre-spotted 384-well compound plates. Each plate included uninfected mock-control wells for normalization. The plates were incubated at 37 °C and 5% CO_2_ atmosphere for 24 h or 48 h, depending on the assay set up.

### Viability screen

The viability screen was conducted using five cell lines: Vero E6, Calu-3, U2OS, HepG2, and Caco-2; and were tested in five-point dose response ranging from 1 nM to 10 μM. Cells were seeded at 25 µl per well to pre-spotted 384-well white cell culture plates (Greiner Cellstar) using Multidrop Combi (Thermo Scientific) and were incubated at standard conditions for approximately 72 h. The plates were placed in humidity boxes to maintain optimal conditions during incubation. Cell viability was assessed using the CellTiter-Glo 2.0 assay (Promega), following the manufacturer’s protocol and luminescence was measured using the Enspire reader (PerkinElmer). The data was analyzed using the web-based software Breeze with Drug Sensitivity Score (DSS) as one of the readouts, a metric derived from dose-response curves (Potdar et al., 2023; Yadav et al., 2014). A DSS of zero indicates no toxicity, while a DSS > 10 indicates compound toxicity.

### Antiviral Rescue of Cytopathic Effect (CPE)

Infected Vero E6 cells were seeded at a density of 4000 cells per well in 30 μL using a Viaflo 384 (Integra Biosciences) and incubated under standard conditions. Compounds were tested in five doses, ranging from 0.001 to 10 μM, with uninfected and infected controls. At 48 hours post-infection (hpi), cell viability was measured by luminescence using the SpectraMax iD5 (Molecular Devices) and the CellTiter-Glo assay 2.0 (Promega), following the manufacturer’s protocol. The data were normalized to uninfected and infected DMSO control wells, and inhibition of cytopathicity (%) was calculated relative to these controls. A compound was defined as a hit if any of the tested doses resulted in an antiviral efficacy exceeding 15%.

### Morphological profiling and immunostaining

Infected Vero E6 cells (2500/well, 30 μL), and A549^ACE2^ cells (4000 cells/well, 30 μL) were seeded using a Viaflo 384 onto pre-spotted 384-well assay plates (Phenoplate, Revvity). The plates were then incubated for 24 h, after which they were fixed with 4% paraformaldehyde (Thermo Scientific), washed twice with PBS, and stained using our adapted Cell Painting assay (Rietdijk et al., 2021). The cells were permeabilized with Triton-X (0.1% Triton X-100), blocked with BSA (Sigma), and incubated overnight with primary antibodies: anti-spike (GeneTexA) for Vero E6 and anti-nucleocapsid (Invitrogen) for A549^ACE2^. Then plates were washed 3x with PBS, followed by staining with a dye mix including Hoechst, SYTO 14, Concanavalin A, WGA, Phalloidin, and a secondary antibody (Invitrogen). Plates were washed an additional 3x with PBS, and then stored at 4 °C until imaged. Liquid handling was automated using an automated reagent dispenser (Biotek Multiflo FX).

Fluorescent images were acquired using an Image Xpress Micro XLS (Molecular Devices) microscope with a 20X objective using laser-based autofocus for Vero E6 experiments, or with a Squid (Cephla) microscope for A549^ACE2^ cells. In total, 9 sites per well were captured using 5 fluorescence channels. The images were analyzed with the open-source image analysis software CellProfiler version 4.2.1 (Stirling et al., 2021). The Cellpose v2.0 plugin, using the cyto2 model was used for segmentation of nuclei and cells (Stringer et al., 2021). A quality control pipeline and illumination correction pipeline was used to remove corrupted images. A total of 2122 features per object were extracted using CellProfiler’s AreaShape, Correlation, Intensity, Granularity, Location, Neighbors, and RadialDistribution modules.

Downstream analysis of the feature data was performed using Python 3, using Jupyter notebooks. Images flagged as outliers based on a threshold of ±10 standard deviations from the mean were excluded. Mean feature values were computed on the well level. Invariant features (standard deviation ≤ 0.0001) or features with extreme variation (>15 standard deviations), and features with missing data were removed. MAD normalization was applied by subtracting the median and dividing by the MAD of the corresponding DMSO samples on each plate. To reduce the influence of extreme values, all feature values were clipped to the range of -50 to 50.

Partial Least Squares (PLS) regression was used to model the phenotypic separation between infected and uninfected controls, where uninfected wells were labeled as 1 and infected wells as 0. Predictive power (*Q*^2^) and accuracy (*R*^2^) were calculated for the PLS-DA model, which were estimated by three-fold cross-validation. For each compound, the mean prediction score was calculated reflecting the extent to which the compound’s phenotypic profile resembled the uninfected state. In parallel, we calculated the Pearson correlation coefficient between each compound’s profile and the mean profile of the uninfected controls. To quantify overall similarity to the uninfected phenotype, we averaged the PLS-DA prediction score and the correlation coefficient, yielding a morphology-to-uninfected-control score (referred to as *morphology score*) for each compound. Processing of features for the analysis of non-infected cell populations was performed using Pycytominer (Serrano et al., 2025).

For visualization of the affected features in radar plots, the absolute mean of the normalized features was computed, grouped by CellProfiler module (Intensity, Correlation, Granularity, Location, RadialDistribution), and stain (Hoechst, Concanavalin A, SYTO 14, Mitotracker, WGA and Phalloidin). Area-shape related features were grouped by cell compartment, i.e. Cell (C), Cytoplasm (Cy) and Nucleus (N). Neighboring related features were grouped by Cell (C) and Nucleus (N) and Neighbour features were grouped per cell compartment, i.e. correlation features between different stains were denoted by their initial letters.

The antibody staining included in the assay, was used to quantify the number of infected cells per well. A cell was considered infected if the virus-specific antibody intensity exceeded a threshold defined as the mean plus three standard deviations of the antibody signal in the blank (uninfected) control wells. To minimize batch effects, the threshold was calculated for each plate.

### Drug-induced phospholipidosis (DIPL) evaluation

Compounds were dispensed in 384-well plates as previously described, in six three-fold dilutions (0.3 - 30 μM) and in technical duplicates. A549^ACE2^ cells were split, spun down and resuspended in serum-free, phenol red-free DMEM/F-12 (Gibco) media. The cells were incubated with CellTracker Deep Red Dye (13 μM, Thermo Fisher Scientific) for 40 min, centrifuged and resuspended with serum-containing phenol red-free DMEM/F-12 media.

Hoechst 33342 (120 nM, Invitrogen) and HCS LipidTOX Green Phospholipidosis Detection Reagent (0.25X, Thermo Fisher Scientific) were added to the cell suspension. The cells were seeded using multidrop (Thermo Fisher Scientific) at a density of 1500 cells per well (30 μL) and were maintained at standard conditions. The plates were kept at standard conditions and imaged live with the Operetta microscope (Revvity), using 20X confocal water objective, at 24 h, 48h, and 72h post-seeding. The images from the Operetta were subsequently analyzed using the Harmony software (Revvity, USA).

An analysis pipeline for detecting and quantifying phospholipidosis was established based on the average area of spots per cell, detected by the LipidTOX dye. Assay quality (0.3% DMSO as negative control and 10 μM tamoxifen as a positive control) was calculated according to standard procedures and plates above Z’-factor threshold of 0.4 were considered for the experiment (Qian, 2025). Defining the non-DIPL inducers, a primary threshold of 9% DIPL was calculated as mean plus three standard deviations of the DMSO control. A secondary threshold of 20% DIPL was set to distinguish the low from moderate-to-high DIPL inducers.

Compounds were fitted to a four-parameter logistic (LL.4) model to create dose-response curves and the curve fitting was performed in R (version 2024.12.0+467) using the drc (v3.0-1) package.

### Pathway enrichment analysis

Gene Ontology (GO) enrichment analysis was performed on the protein targets annotated in Reinshagen et al., sourced from ChEMBL bioactive compound targets of the 74 validated compounds (Reinshagen et al., 2025). *Homo sapiens* proteins with a confidence score ≥ 8 and pChEMBL score ≥ 6 from binding and functional assays were included. The analysis was conducted using the *enrichGO()* function from the *clusterProfiler* R package (v4.14.6), focusing on the Biological Process ontology and the human gene annotation database *org.Hs.eg.db*. The following parameters were applied; p-value adjustment by the Benjamini-Hochberg method, p-value cutoff of 0.05, and a q-value cutoff of 0.2. The enrichment was visualized using the top 100 GO terms using *enrichmentNetwork()* function from the Advanced Pathway Enrichment Analysis Representation (*aPEAR)* package (v1.0) (Kerseviciute and Gordevicius, 2023)).

### Reagents and tools table

| Reagent/Resource | Reference or Source | Identifier or Catalog Number |
| --- | --- | --- |
| **Experimental models** |  |  |
| Vero E6 (*C. sabaues*) | Donated by cell profiling unit<br>Charlotte Stadler |  |
| Calu-3 (*H. sapiens*) | ATCC | HTB-55 |
| U2OS (*H. sapiens*) | Donated by cell profiling unit<br>Charlotte Stadler |  |
| HepG2 (*H. sapiens*) | Donated by cell profiling unit<br>Charlotte Stadler |  |
| Caco-2 (*H. sapiens*) | Donated by Jordi Carreras<br>Puigvert |  |
| A549<sup>ACE2</sup> (*H. sapiens*) | Donated by Oscar<br>Fernandez-Capetillo lab | Porebski et al (2024)<br>(Molecular Therapy) |
| U87-MG (*H. sapiens*) | ECACC | 890814 |
| Vero (*C. sabaues*) | ATCC | CCL-81 |
| MRC-5 | ATCC | CCL-171 |
| CoV-2/human/SWE/01/2020 | Lentini, A., Papanicolaou, N.,<br>& Reinius, B. (2021) | GenBank: MT093571.1 |
| HCoV-229E | ATCC | VR-740 |
| Zika virus | European Virus Archive | Strain H / PF / 2013 |
| **Antibodies and stains** |  |  |
| Mouse monoclonal Anti-spike<br>antibody | GeneTex | GTX632604 |
| Mouse monoclonal<br>anti-nucleocapsid protein | Invitrogen | B46F |
| Goat anti-mouse IgG | Invitrogen | A-21235 |
| Hoechst 33342 | Invitrogen | H3570 |
| SYTO 14 | Invitrogen | S7576 |
| Concanavalin A | Invitrogen | C11252 |
| Phalloidin | Invitrogen | 10135092 |
| CellTracker Deep Red Dye | Invitrogen | C34565 |
| HCS LipidTOX Green<br>Phospholipidosis Detection<br>Reagent | Invitrogen | H34350 |
| **Chemicals, Enzymes and**<br>**other reagents** |  |  |
| SPECs drug repurposing<br>library | https://www.specs.net/ |  |
| DMEM High-glucose | Gibco | 11965092 |
| L-glutamine | Cytiva | SH30034.01 |
| DMEM/F-12, GlutaMAX Supplement | Gibco | 11514436 |
| Non-essential amino acids | Cytiva | SH30238.01 |
| McCoy's 5A medium | Sigma-Aldrich | M8403 |
| MEM with Earl's salts | Sigma-Aldrich | M2279 |
| Heat-inactivated FBS | Gibco | 10500064 |
| Penicillin/streptomycin | Cytiva | SV30010 |
| D-PBS | Nordic biolabs | SH30028.FS |
| TrypLE | Gibco | 11538856 |
| CellTiter-Glo | Promega | G9243 |
| Paraformaldehyde | Thermo Scientific | 28908 |
| BSA | Sigma | A8022 |
| **Software** |  |  |
| CellProfiler (v.4.2.1) |  |  |
| R package *clusterProfiler* (v4.14.6) | G Yu (2024) (The Innovation) |  |
| R package *aPEAR* (v1.0) | Kerseviciute I, Gordevicius J. (2023) (bioRXIV) |  |
| R package drc (v.3.0-1) | Ritz C et al (2015) (PLOS ONE) |  |
| GraphPad Prism (v.10.5.0) | https://www.graphpad.com/ |  |
| Harmony | Revvity |  |
| **Other** |  |  |
| MycoAlert mycoplasma detection kits | Lonza |  |
| Celphia Squid microscope | Cephla |  |
| Image Xpress Micro XLS | Molecular Devices |  |
| Operetta | Revvity |  |

## 3. Results

### 3.1. Morphological profiling captures virus-induced effects and enables screening for antiviral compounds

To demonstrate the utility of morphological profiling for antiviral compound screening against SARS-CoV-2, we employed our previously established morphological profiling approach, combining antibody-based detection of viral proteins with Cell Painting-based morphological analysis (Rietdijk et al., 2021). This combined assay enables the characterization of virus-induced morphological changes and screening for compounds capable of reversing the virus-induced phenotype.

The workflow begins with seeding pre-infected cells into multiwell plates containing compounds. Following a 24-hour incubation period, cells are fixed and subjected to multiplexed (immuno-)fluorescence staining. This includes antibody-based labeling of viral proteins (e.g., nucleocapsid or spike proteins) alongside Cell Painting stains targeting major cellular components such as the nucleus, nucleoli, endoplasmic reticulum, Golgi apparatus, plasma membrane, and cytoskeleton. High-throughput imaging is performed to capture multiple sites per well, after which the raw images are processed through an image analysis pipeline that segments individual cells and extracts a large set of morphological features (>1,500 per cell) describing intensity, texture, and shape. These feature vectors are then mean aggregated and normalized to generate well-level morphological fingerprints, which are analyzed to identify compounds that mitigate or revert the virus-induced phenotype (**Fig. 1A**).

**Figure 1.**
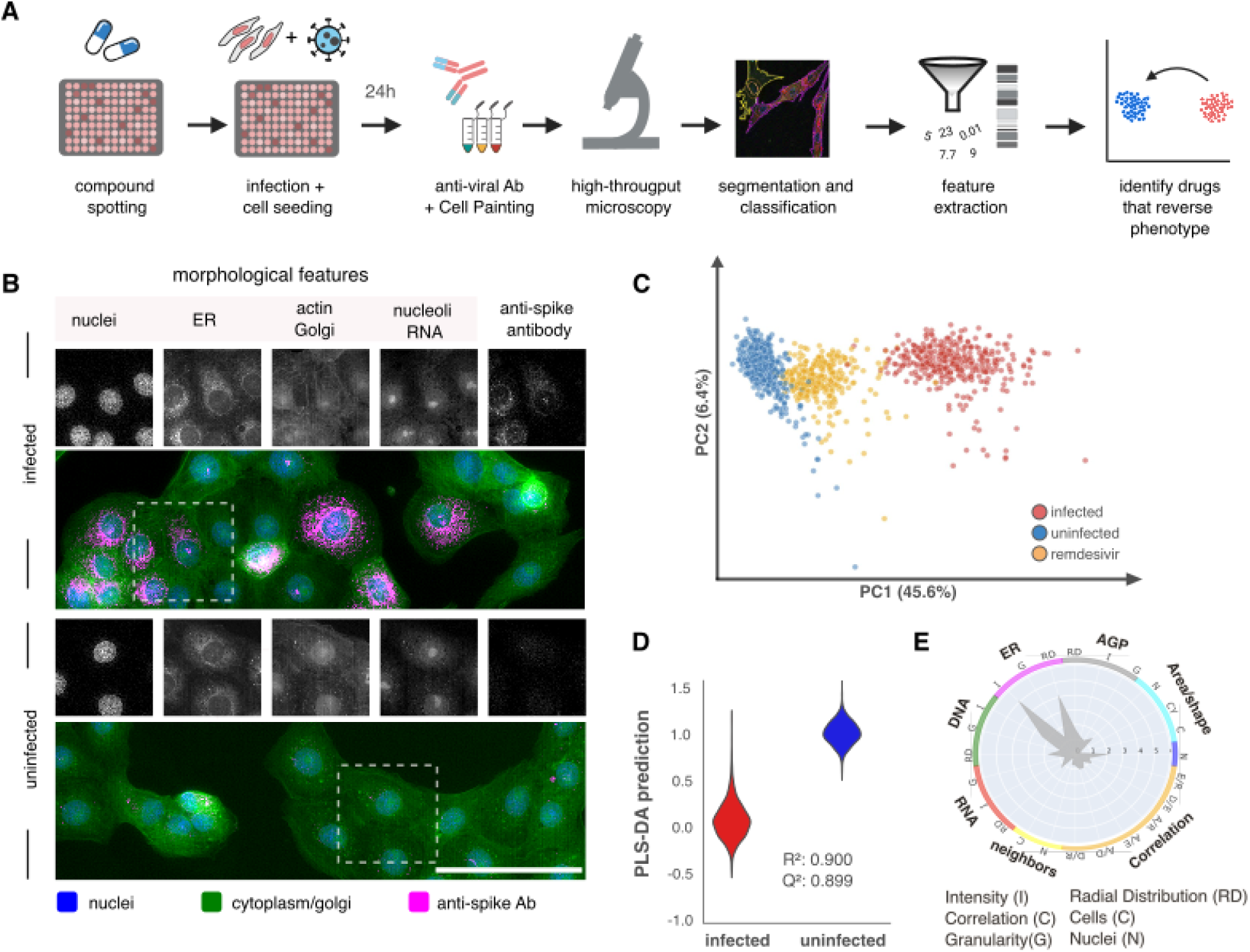
Morphological Cell Profiling workflow adapted for antiviral drug discovery. **A.** Schematic overview of the experimental workflow. Compounds are pre-spotted onto assay plates, followed by seeding of pre-infected cells. After 24 hours of incubation, cells are fixed and stained with both a virus-specific antibody and Cell Painting dyes. This is followed by high-throughput imaging and feature extraction, enabling characterization of a virus induced phenotype and screening of compounds that could reverse the virus-induced phenotype. **B.** Representative images of Vero E6 cells, either infected with SARS-CoV-2 or uninfected controls for each of the five imaging channels. **C.** Principal Component Analysis (PCA) applied to morphological features, with each point representing a single well. Percent variance explained by each component is indicated. **D.** Observed versus predicted values from the PLS-DA (Partial Least Squares Discriminant Analysis) model trained to classify infection status based on cell morphology features and model performance metrics. **E.** Radar plot illustrating the most significantly changed morphological feature groups in response to viral infection. Scale bar = 100 μm

**Figure 1B** shows representative images from Vero E6 cells infected with SARS-CoV-2 across the five fluorescent channels. Approximately 70% of the cells are infected, as indicated by the antibody staining. To assess whether infection induces detectable morphological changes, we performed principal component analysis (PCA) on the extracted morphological features (excluding antibody signals). As shown in **Figure 1C**, PCA revealed clear separation between infected, uninfected, and remdesivir-treated profiles, indicating a distinct infection-induced phenotype that is partially rescued by antiviral treatment. Supervised classification using partial least squares discriminant analysis (PLS-DA) yielded R² and Q² values of 0.97, indicating strong separation between infected and uninfected samples based on morphological features alone (**Fig. 1D**).

The most pronounced alterations upon infection are the endoplasmic reticulum (ER) intensity and radial distribution features (**Fig. 1E**). This aligns with the images shown in **Fig. 1B**, which displays increased ER brightness in infected cells. Additionally, the number of nuclei per cell is affected, potentially indicating the formation of multinucleated cells, known as syncytia.

### 3.2. Drug repurposing screens against SARS-CoV-2

To identify repurposable drug candidates against SARS-CoV-2, we systematically screened a library of 5,275 compounds using three complementary assays: a cell viability assay to assess compound toxicity, an antiviral rescue of cytopathic effect (CPE) assay, and the combined morphological profiling with antibody staining. The compound library, originally curated by the Drug Repurposing Hub, comprises a chemically diverse set of small molecules that have reached clinical development stages, including approved, trial-phase, and withdrawn drugs. Most compounds have known mechanisms of action and target classes such as G-protein-coupled receptors, kinases, ion channels, nuclear hormone receptors, and voltage-gated sodium channels (Corsello et al., 2017).

#### 3.2.1 Cell viability screen

When screening a large and chemically diverse compound library, varying compound potencies can result in missed compound activity if only a single concentration is tested, especially when cytotoxicity masks antiviral efficacy. To address this, we first performed a viability screen to assess compound toxicity. Viability was evaluated across five cell lines using the drug sensitivity score (DSS), allowing a generalizable assessment of cytotoxic effects (**Fig. 2A, supplementary file 1-5**). Overall, the drug repurposing set caused no major signs of toxicity across the cell lines, with an average DSS of 1.3. HepG2 and Calu-3 cells exhibited slightly higher compound toxicity, with 5.7% and 4.9% of compounds with a DSS >10, respectively. Based on the viability screen, the highest non-toxic concentration, either 0.83 or 10 µM, with >80% viability in Vero E6 cells was selected for further experimentation. This concentration also guided dose selection for the subsequent validation screens, using a range spanning below and above the hit concentration.

**Figure 2.**
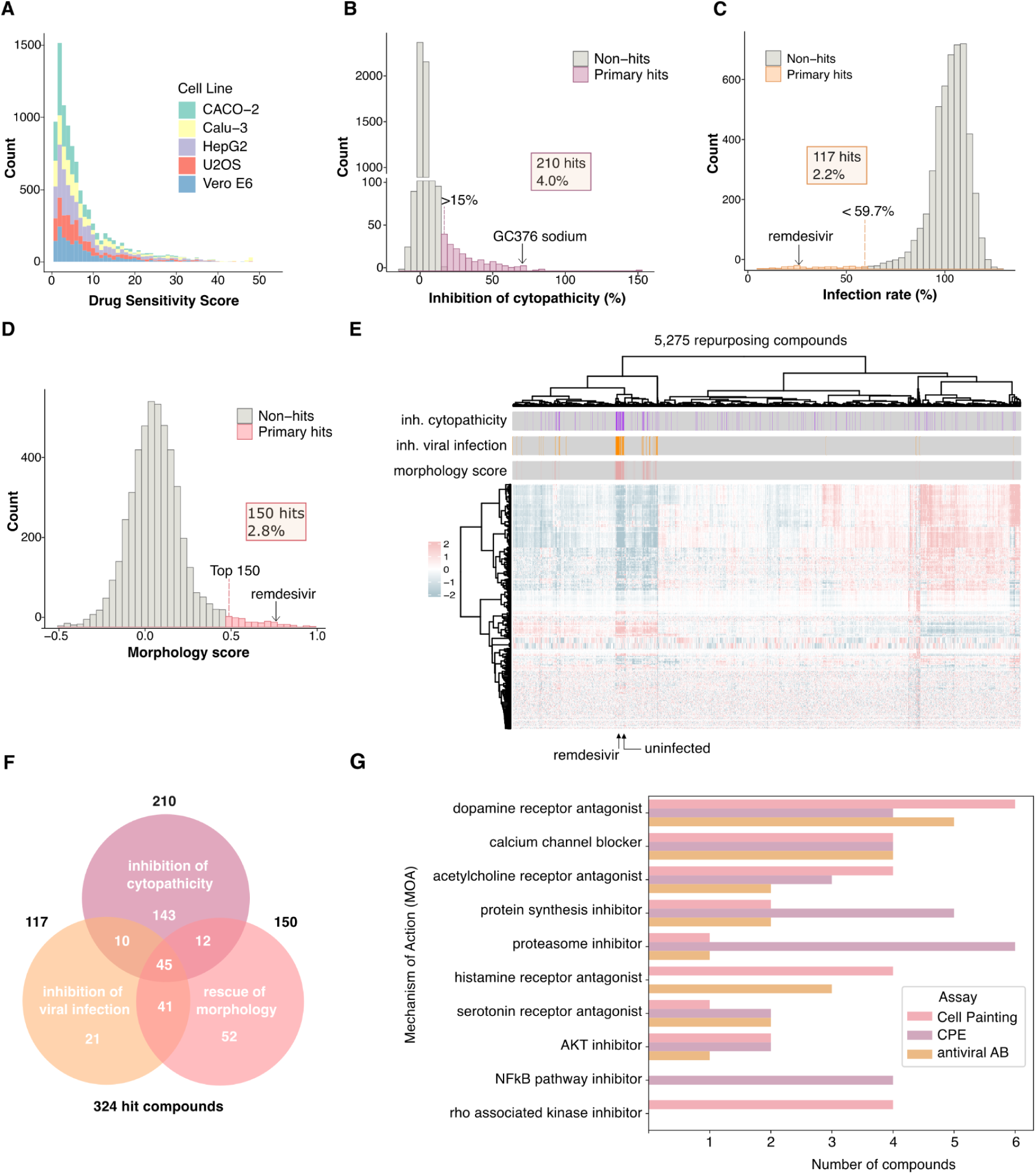
Screening of a drug repurposing library of 5,275 compounds using four assay readouts. **A.** Overview of viability screening across five cell lines, used to calculate sensitivity scores and inform concentration selection for follow-up assays. **B–E** Histograms showing the distribution of assay values for: **B.** cytopathicity inhibition (best dose selected per compound), **C.** Viral infection rate, and **D.** morphology scores. Thresholds used to define hits are indicated. **E.** Unsupervised hierarchical clustering of morphological profiles for all 5,275 compounds using Ward’s linkage method and the Euclidean distance metric; on top of the heatmap we show the hits for the three respective read-outs. **F.** Venn diagram depicting the overlap of compounds identified as hits across the three assay readouts. **G.** Bar chart summarizing the number of compounds corresponding to the top 10 mechanisms of action, with counts shown per assay type.

#### 3.2.2 Antiviral rescue of Cytopathic Effect (CPE)

In the CPE assay, compounds were evaluated for their ability to prevent virus-induced cell death, serving as a functional readout of phenotypic antiviral activity. The full compound library was screened across five concentrations from 1 nm to 10 µM. A small subset of compounds reduced virus-induced cytopathicity (**Fig. 2B, supplementary file 6)**. We selected compounds that, at one or more tested concentrations, mitigated the virus-induced reduction in cell viability by 15% or more, resulting in a set of 210 candidate compounds.

#### 3.2.3 Antibody-based screen

Antibody detection provides a direct approach for identifying compounds that reduce viral infectivity. Here we used the antibody channel, which was included in the morphological profiling assay, to identify compounds that lowered viral infection levels. Compounds that reduced infection by more than 40% were selected, yielding a total of 117 candidate compounds (**Fig 2C, supplementary file 7**).

#### 3.2.4 Morphological profiling

To identify compounds capable of reversing the virus-induced morphological phenotype, we applied a morphology-based scoring approach that quantified each compound’s similarity to the uninfected control profile. Compounds with higher scores exhibited profiles more closely resembling non-infected controls. Based on this metric, we prioritized the top 150 compounds for follow-up experimentation (**Fig 2D, supplementary file 7**). To further explore phenotypic similarities among these hits, we applied hierarchical clustering with Ward’s linkage and Euclidean distance. This analysis revealed a cluster of compounds that shared similarity in terms of their morphology to the non-infected controls. This cluster was enriched for compounds that reduced the number of infected cells and inhibited cytopathic effects (**Fig. 2E**).

Based on the results from these three primary screening read-outs, we selected the hits from each of read-outs for further validation. In total, 324 compounds were prioritized for follow-up testing. Among these, there was an overlap of 12 compounds between the Cell Painting and CPE assay, 41 compounds between the Cell Painting and antibody-based screen, and 10 compounds between the antibody-based screens and the CPE assay. A total of 45 compounds were identified as hits across all three assay types (**Fig 2F**). We found that the majority of compounds which reduced viral infectivity were also associated with a shift toward uninfected morphology. In contrast, compounds that prevented virus-induced cell death (cytopathic effects) did not consistently correlate with a rescue of morphology (**Fig EV1A**).

The three assays identified compounds with diverse mechanisms of action (MoA), covering a total of 204 distinct mechanisms. **Figure 2G** shows the top ten mechanisms of action (MoAs) among the hit compounds, along with the number of compounds identified per assay for each MoA. The most prevalent mechanism of action (MoA) among the hit compounds was dopamine receptor antagonist, which was identified in all three assays. Notably, morphological analysis uniquely identified Rho-associated kinase (ROCK) inhibitors as hits that were not detected in the other two readouts. Conversely, inhibitors of the NF-κB pathway were exclusively identified in the CPE assay.

### 3.3. Hit validation in human lung epithelial A549^ACE2^ cells

To validate the compounds identified in the three primary screens, we tested the 324 hit compounds in a four-point dose-response assay (0.1-3.3 µM or 0.3-30 µM) in human A549^ACE2^ cells using morphological profiling. To confirm that A549^ACE2^ is a suitable cell model, we first evaluated whether SARS-CoV-2 infection induced a detectable morphological phenotype in this cell model. As shown in **Figure 3A**, representative images of infected and non-infected control wells revealed clear morphological changes across the different channels. Using PLS-DA, we found that morphological features alone could effectively separate infected from non-infected conditions, achieving an area under the ROC curve (AUC) of 1.000, and a *R*^2^ and *Q*^2^ score of 0.973 and 0.969 respectively (**Figure 3B**). Phenotypic feature groups that changed significantly upon infection included endoplasmic reticulum intensity, as well as alterations in cytoskeletal stains and Golgi and correlation-based features (**Figure 3C**). A total of 208 (64.8%) compounds exhibited the ability to reverse the virus-induced phenotype (morphology score >0.5). We observed dose-dependent rescue of cellular morphology following treatment of effective antiviral compounds (**Figure 3D, Supplementary file 8**). When comparing responses between the two cell lines, compound effects varied considerably, and no strong correlation was observed between the morphological scores in Vero E6 and A549^ACE2^ cells (**Fig EV1B**). Nevertheless, 81 (25%) of the tested compounds achieved a morphology score above 0.5 in both cell lines, indicating partial to substantial reversal of the virus-induced phenotype (**Fig EV1C, Supplementary file 8**).

**Figure 3.**
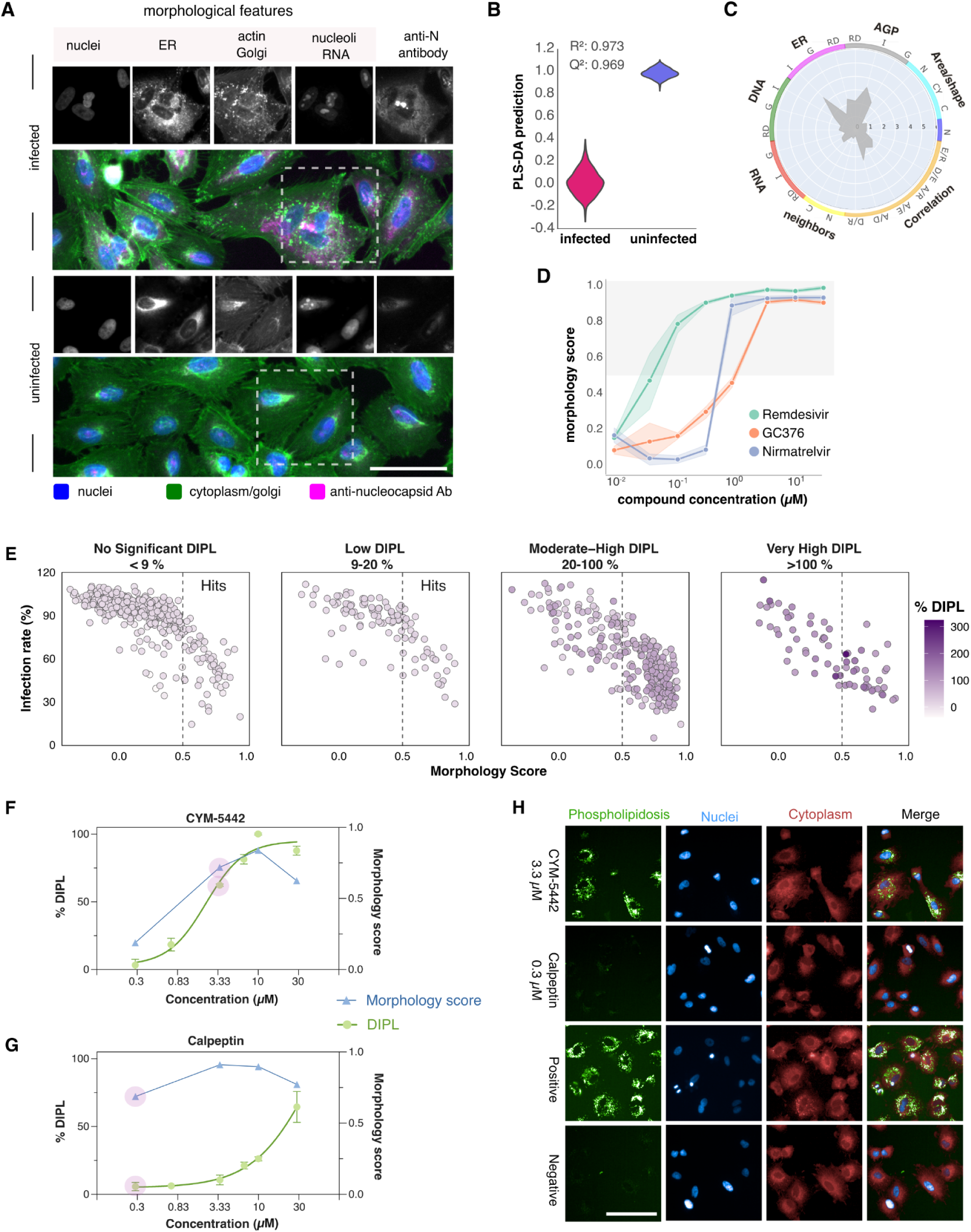
Validation and deprioritization of antiviral compound hits in human lung epithelial cells using morphological profiling and DIPL assessment. **A**. Representative Cell Painting images of infected and non-infected A549^ACE2^ control wells showing distinct morphological changes upon SARS-CoV-2 infection. Scale bar = 100 μm. **B.** Observed versus predicted values from the PLS-DA (Partial Least Squares Discriminant Analysis) model trained to classify infection status based on cell morphology features and model performance metrics. **C**. Radar plot illustrating the most significantly changed morphological feature groups in response to viral infection. **D.** Dose–response plot of antiviral control compounds based on morphology scores. Shaded regions show the 95% confidence interval. The grey area highlights the dose range where compounds successfully reversed the virus-induced phenotype. **E.** Faceted scatter plots showing the relationship between morphology score (x-axis) and SARS-CoV-2 infection rate (y-axis), stratified by drug-induced phospholipidosis levels (% DIPL). Each panel represents a DIPL category and each point represents a compound-concentration pair from the 324 hit compounds, colored by % DIPL intensity. The vertical dashed line indicates the morphology score threshold (>0.5). Selected hits fall within the range ≤20% DIPL and morphology score >0.5. **F-G.** Dose-response curves of CYM-5442 and calpeptin, showing %DIPL (left y-axis, green color) and PLS-correlation (right y-axis, blue color), across log-transformed compound concentrations (x-axis). The lowest concentration reaching the morphology score threshold (>0.5) is annotated in purple. DIPL data points are represented by the mean ± SD of two replicates and dose-response curves were fitted using a four-parameter model in GraphPad Prism (v.10.5.9) **H**. Representative fluorescence images of uninfected A549^ACE2^ cells treated with CYM-5442 (3.3 μM), calpeptin (0.3 μM), positive control (tamoxifen) and negative control (DMSO), at 24h. Stains: phospholipidosis (green), nuclei (blue), cytoplasm (red). Scale bar = 100 μm.

To prioritize antiviral candidates, we integrated the morphology scores, infection rates and drug-induced phospholipidosis (DIPL) responses (**Fig. 3E**). Compounds were classified as no significant DIPL (<9%), low (9-20%), moderate-high (20-100%), and very high (>100%). A trend of decreased infection rate with increasing DIPL levels was observed, this was especially apparent beyond 20% DIPL (**Fig. 3E, fig EV2A**). Based on this, a threshold of deprioritizing compounds with >20% DIPL was set. A total of 74 compounds were selected that had a morphological score (>0.5) and no to moderate DIPL levels (≤20%). The selection process is further visualized in **Fig EV2B**. In **Figure 3F-G**, we illustrate the relationship between the DIPL score and the morphology score for a deprioritized compound (**Fig. 3F**), as well as a prioritized hit compound (**Fig. 3G**). While both compounds demonstrate a phenotypic rescue (morphology score >0.5), CYM-5442 exceeds the DIPL threshold at its active antiviral concentration and was therefore deprioritized (**Fig. 3F**). In contrast, calpeptin induced morphological rescue without exceeding DIPL limits at its lowest effective antiviral dose (**Fig. 3G**). Representative immunofluorescent images of both compounds show the phospholipid accumulation in CYM-5442-treated cells, comparable to the positive control (tamoxifen); and the absence of such accumulation in calpeptin-treated cells, comparable to the negative (DMSO) control (**Fig. 3H**).

In addition, a clear dose-dependency in phospholipidosis was observed. At 0.3 and 0.8 μM, most compounds (89.4% and 73.2%) exhibited no significant DIPL, while concentrations from 3.3 μM and above showed higher levels of DIPL; at a concentration of 30 μM, 50.2% of the compounds induced moderate-high DIPL (**Table 1, Supplementary file 9**). Moreover, we evaluated the DIPL development across three timepoints (24h, 48h, 72h) and observed that EC_50_ values were lowest at 24 h, indicating the most sensitive timepoint for phospholipidosis detection, with EC_50_ values reached at lower concentrations (**Fig EV2C, Supplementary file 9-12**). For this study, the 24 h timepoint was evaluated to align with the time of infection. Additionally, five impure compounds and previously reported DIPL inducers not active in our screen due to low solubility or cell line specificity were excluded from further analysis.

**Table 1.** Distribution of compounds across concentrations and DIPL categories.

| DIPL category / Concentration (μM) | 0.3 | 0.8 | 3.3 | 6.6 | 10 | 30 |
| --- | --- | --- | --- | --- | --- | --- |
| No significant DIPL (<9%) | 89.4% | 73.2% | 45.1% | 34.4% | 31.2% | 23.9% |
| Low DIPL (9-20%) | 8.4% | 16.3% | 12.8% | 10.7% | 9.2% | 9.0% |
| Moderate–High DIPL (20-100%) | 1.6% | 10.2% | 37.5% | 41.6% | 41.4% | 50.2% |
| Very High DIPL (>100%) | 0.6% | 0.3% | 4.6% | 13.5% | 18.2% | 16.9% |

### 3.4 Integrated target-pathway enrichment and hit prioritization reveal repurposed compounds with possible host-targeting activity

To gain insight into the 74 validated compounds, we explored their annotated targets (n = 649), which had a representation across eight different protein classes: enzymes, membrane receptors, transporters, transcription factors, ion channels, epigenetic regulators, cytosolic proteins, and G protein-coupled receptors (GPCRs) (**Fig. 4A**) (Reinshagen et al., 2025). The most prominent protein class was enzymes, with kinases representing the most frequently annotated subclass.

**Figure 4.**
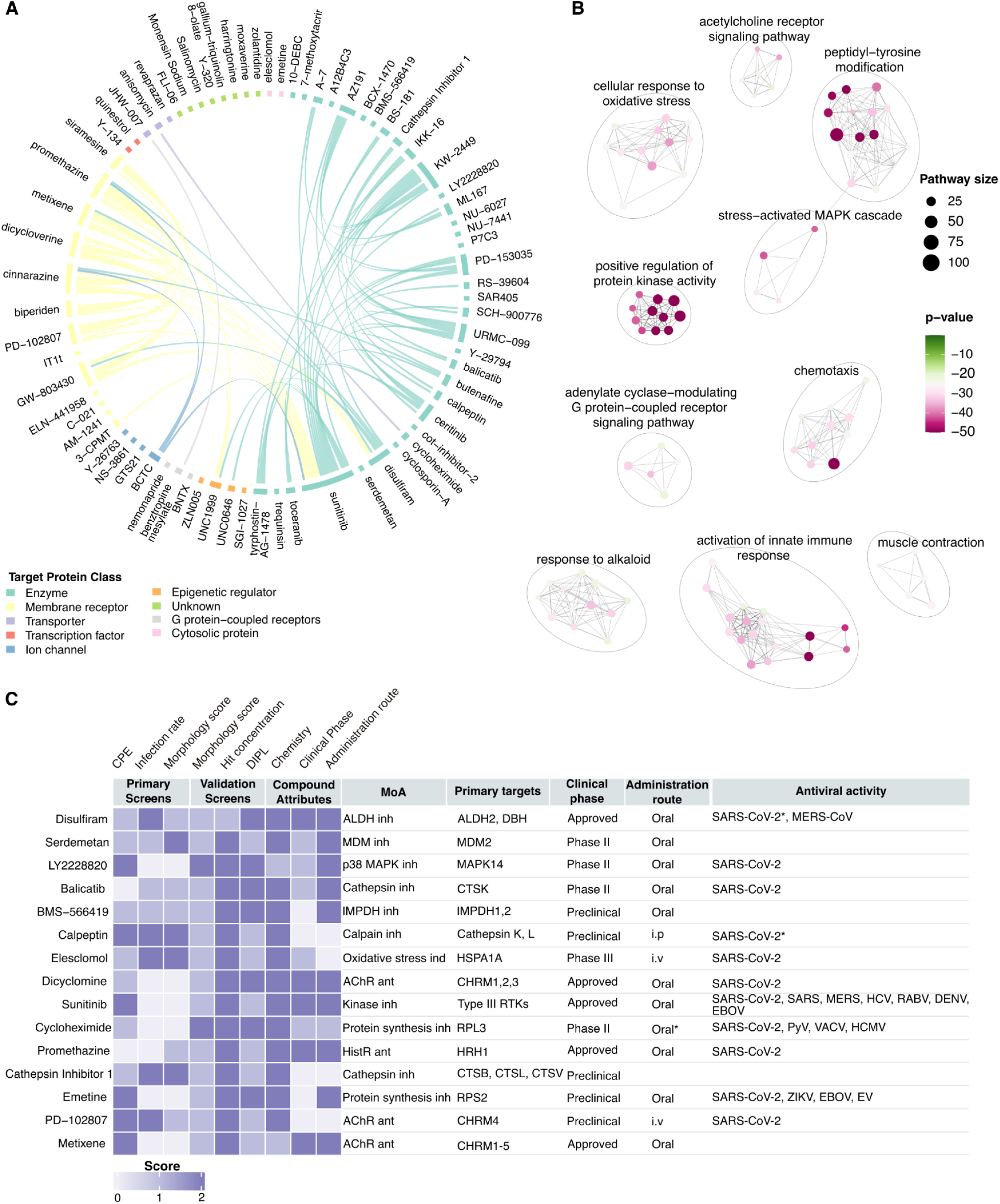
Host biological pathways across the protein targets and prioritization of antiviral hits. **A**. Chord diagram of the host protein target overlapping the 74 validated compounds. The compounds are grouped by target protein class, indicated by the color and lines connecting compounds with shared targets. **B**. Gene Ontology (GO) enrichment analysis on the 649 annotated targets visualized by the top 100 enriched terms (adjusted p-value < 0.05). Nodes represent the pathways, and the edges represent the similarity between the pathways. Node size represents the pathway size and is colored by p-value (-log10). **C**. Hit scoring system towards pandemic preparedness by integration of screening data with compound attributes. A heatmap of the top 15 ranked repurposable compounds based on combined scores from the primary screens, validation screens, and compound attributes. The chemistry category includes molecular weight (MW) and logP value. MW ≤500 and logP≤5 was ranked highest. The hits were given points 0-2 in each category and were summarized for ranking and listed in descending order. Additional metadata includes MoA, primary annotated targets from the Clue.io Repurposing Hub (Corsello et al., 2017), highest clinical phase, administration route, and reported SARS-CoV-2 and other viral activity. Oral* indicates the compound is only used experimentally. SARS-CoV-2* indicates activity against viral proteins. CPE - Cytopathic effect assay, DIPL - drug-induced phospholipidosis, inh - inhibitor, ind - inducer, ant - antagonist, i.p - intraperitoneal, i.v - intravenous, MERS-CoV - Middle East respiratory syndrome-related coronavirus, SARS-CoV - severe acute respiratory syndrome coronavirus, HCV - hepatitis C virus, RABV - rabies virus, DENV - dengue virus, EBOV - ebola virus, PyV - polyomavirus, VACV - vaccinia virus, HCMV - human cytomegalovirus, ZIKV - zika virus, EV - enterovirus.

We next assessed target overlap across the compounds by grouping them according to their predominant protein class and linking compounds that share targets (**Fig. 4A**). In total, 182 shared targets were identified. To further explore the biological pathways overrepresented among the targets, we conducted a GO enrichment analysis and visualized the 100 most significantly enriched biological pathways (**Fig. 4B**). Prominent clusters were positive regulation of protein kinase activity, activation of innate immune response, cellular response to oxidative stress, and peptidyl-tyrosine modification associated with stress-activated MAPK signaling.

To characterize compounds with favorable properties for advancement as antiviral drugs, we developed a hit scoring system that integrated performance in the primary and validation screens with a TPP, which defines the main desired attributes suitable for a drug against COVID-19. These included chemical profile (molecular weight and logP value), clinical phase, and route of administration, where more clinically advanced and orally available compounds were given highest scores. The 15 top-ranking compounds are shown in **Figure 4C** (**Supplementary file 14**). The panel includes compounds across clinical stages, of which 10 are orally formulated. Several compounds share MoA and targets, including three acetylcholine receptor (AChR) antagonists, dicyclomine hydrochloride, PD-102807, and metixene; and three cathepsin inhibitors, balicatib, calpeptin and cathepsin inhibitor 1.

Of the top 15 hit compounds, 11 have previously demonstrated antiviral activity against SARS-CoV-2, of which two are directly targeting SARS-CoV-2 proteins. Furthermore, sunitinib, cycloheximide, and emetine dihydrochloride, exhibit antiviral activity towards several other viruses beyond SARS-CoV-2 and the Coronaviridae family. The three highest ranking compounds were disulfiram, serdemetan, and LY2228820.

### 3.5 Morphological profiling provides rich profiles that can be used for off-target effect detection and mechanistic insights

To take full advantage of the high-dimensional morphological profiles, beyond solely computing a well-level morphology score, we explored their potential to reveal off-target effects and provide mechanistic insights into compound activity.

Leveraging the single-cell resolution of our assay, we split the data into two subsets based on the antibody staining. For each well, all non-infected and all infected cells were aggregated to create a perturbation level morphological profile (**Fig. 5A**). We first evaluated how many compounds resulted in significant morphological effects by calculating mean average precision (mAP) scores at the perturbation level (defined as each unique compound–concentration combination) for non-infected cells. We used cosine similarity to rank all profiles by similarity and determined the statistical significance of the mAP scores using permutation testing (Kalinin et al., 2025). In total, 93% of the compounds exhibited statistically significant (p < 0.05 for corrected p-value) and reproducible morphological effects for at least one of the tested doses. Across all conditions in the screen, 58% of the perturbations induced a significant morphological effect in the uninfected cell populations (**Fig EV3A**).

**Figure 5.**
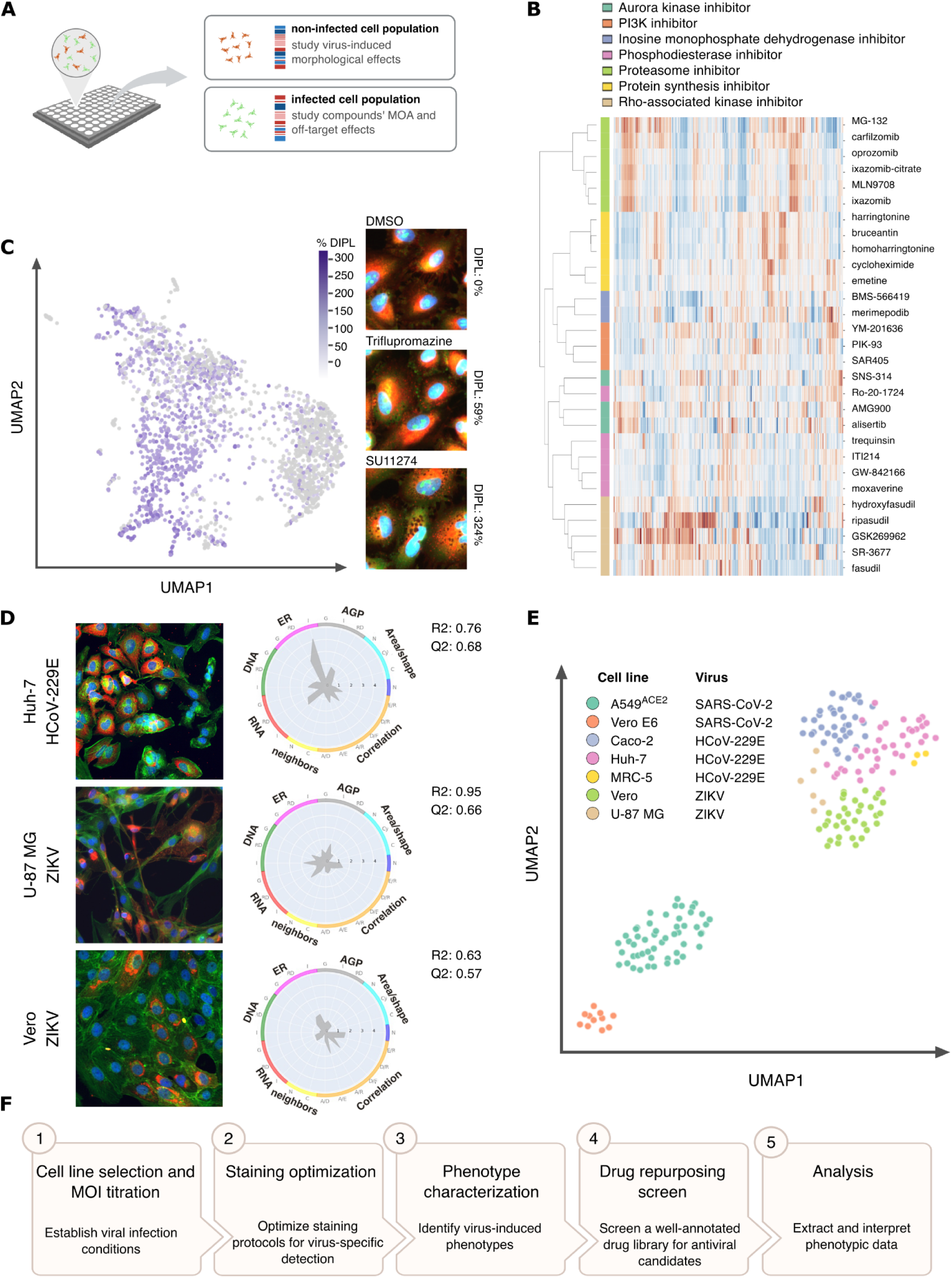
Profiling of infected and non-infected cells reveals virus-specific phenotypes and compound effects. **A**. Schematic overview of the data-splitting strategy: morphological profiles were separated into infected and non-infected cell subsets to enable independent analysis of virus-induced phenotypes and compound-induced mechanisms of action. **B**. Unsupervised hierarchical clustering of compounds based on morphological profiles of non-infected cells, grouped by their annotated mechanisms of action (MoA), using cosine similarity and complete linkage. **C**. UMAP projection of non-infected cell profiles colored by phospholipidosis levels as measured by the DIPL assay. Representative images of selected compounds show characteristic vacuolization for DIPL inducing compounds (blue: nuclei, red: endoplasmic reticulum, green: nucleoli, cytoplasmic RNA). **D**. Representative images and radar plots for virus-infected cells in different cell lines (Huh7 with HCoV-229E, U-87 MG with ZIKV, and Vero with ZIKV). Strength and reproducibility of the profiles are indicated by the R² and Q² values. Fluorescent images labeling used in imaging: nuclei (blue), virus-specific antibody (red), and a cytoplasmic marker including cytoskeleton, Golgi, and plasma membrane (green). **E**. UMAP visualization of morphological profiles of infected cells across different cell lines and viruses **F**. Drug Repurposing Screen for rapid morphological profiling in response to emerging viral outbreaks and pandemic preparedness.

To evaluate whether the morphological profiles could reveal mechanistic insights, we tested whether compounds with the same mechanism of action (MoA) produced similar phenotypic signatures. Using mAP, we selected phenotypically consistent MoAs represented by two or more compounds. Unsupervised hierarchical clustering of compounds from seven selected MoAs, showed that compounds with these same MoAs had similar profiles and clustered together (**Fig. 5B**). Among the compound groups, proteasome inhibitors formed the most distinct cluster, characterized by highly consistent morphological profiles evident in the heatmap. Except for Ro-20-1724, all compounds clustered with other compounds sharing the same MoA.

Next, we assessed whether compounds that induce drug-induced phospholipidosis (DIPL) exhibit distinct morphological profiles. Using the non-infected cell profiles, we generated a UMAP embedding of all conditions with matched doses that had been tested in the phospholipidosis assay (**Fig. 5C**), coloring each point according to the assay results. This revealed that most of the phospholipidosis-inducing and non-inducing compounds cluster separately in morphological space. Representative images show that phospholipid accumulation is visible in Cell Painting images as small dark vesicles in the cytoplasm. A higher number of these vesicles corresponds to compounds that induce greater levels of phospholipidosis.

To further quantify these effects, we calculated the Pearson correlation of all morphological profiles to fluoxetine hydrochloride, a well-characterized phospholipidosis inducer included in the assay. Notably, all compounds with high similarity to fluoxetine hydrochloride (Pearson correlation between 0.7 and 1.0) were identified as moderate or strong DIPL inducers (defined as >20% induction; **Fig EV3B**). A correlation matrix of all compounds (**Fig EV3C**) confirms that most DIPL-inducing compounds exhibit high similarity to one another, including other well-known cationic amphiphilic drugs (e.g. prochlorperazine, triflupromazine). However, not all DIPL-inducing compounds were highly correlated to each other; several compounds with moderate to high DIPL levels showed lower similarity, possibly due to additional compound-induced effects beyond phospholipidosis.

### 3.6 Virus-specific morphological signatures across diverse cell models

To evaluate the broad applicability of our approach across different cell models and viral infections, we profiled three viruses, human coronavirus 229E (HCoV-229E), Zika virus (ZIKV), and SARS-CoV-2, each with two to three permissive cell lines. For every condition, we analyzed the morphological profiles of infected cell populations within each well and normalized them against non-infected cells to reduce well-level confounding effects (**Fig. 5D**).

A UMAP visualization of the resulting morphological profiles (**Fig. 5E**) shows that each virus elicits a distinct and reproducible phenotypic signature. Infected cells tend to cluster according to the infecting virus, regardless of the cell line used, highlighting that virus-specific morphological signatures are conserved across cellular models.

The strength of these morphological effects varies both between viruses and across cell lines, as indicated by the grit scores and Partial Least Squares (PLS) metrics. Among the viruses tested, SARS-CoV-2 consistently induces the most pronounced morphological effects, followed by HCoV-229E and lastly ZIKV. Notably, HCoV-229E–infected cells exhibit specific changes in endoplasmic reticulum morphology, similar to those observed in SARS-CoV-2 infection. In contrast, ZIKV infection results in a distinct morphological signature characterized by various affected correlation features as well as effects on neighboring nuclei, possibly suggesting nuclear aggregation.

### 3.7 Standardized Screening Workflow for Repurposed Antivirals and Pandemic Preparedness

To facilitate rapid screening in the context of emerging viral outbreaks and for pandemic preparedness, we established a set of guidelines and best practices for applying morphological profiling to new cell models and viruses. The recommended workflow consists of five key steps: (1) cell line and multiplicity of infection (MOI) optimization, (2) staining protocol optimization, (3) phenotype characterization, (4) drug repurposing library screening, and (5) data analysis (**Fig. 5F**).

#### Step 2) Staining Optimization

To monitor viral infectivity and ensure consistent infection levels across plates and batches, we recommend incorporating a virus-specific antibody into the staining panel, if available. Since standard Cell Painting uses all five fluorescence channels, one marker must be replaced to accommodate the antibody when using a regular five-channel microscope. We chose removing MitoTracker, as it requires live-cell staining which is less suitable under BSL-3 and 4 conditions. To minimize spectral overlap, we recommend placing the virus-specific antibody in the channel with the highest wavelength (e.g., Cy5/Alexa 647), as antibody signals are often weaker than the Cell Painting dyes. Additionally, careful optimization of washing steps is essential to preserve the cell morphology and maintain high image quality. Disruption of the cell monolayer, such as ruffled or detached cell membranes, can interfere with downstream morphological analysis of the cells.

#### Step 3) Phenotype Characterization

To rapidly extract morphological features from images, we developed an automated CellProfiler pipeline that integrates Cellpose for robust cell segmentation. This pipeline enables high-throughput and consistent analysis of cellular morphology across experimental conditions with no or minor adjustments for various cell lines. By separately identifying nuclei and cytoplasm objects, this pipeline effectively accommodates highly multinucleated cells which is a frequent phenotype observed in virus-infected cells. The first step in phenotype characterization is to evaluate whether viral infection induces a screenable phenotype. This can be assessed by comparing infected versus non-infected conditions. Using the provided Jupyter notebooks, users can load the features from the image analysis pipeline and apply Partial Least Squares Discriminant Analysis (PLS-DA) to determine whether the virus induces a reproducible and distinct morphological signature. We recommend selecting the condition that yields the highest accuracy (R²) and predictive power (Q²) in distinguishing infected from non-infected cells. These metrics will help guide the final choice of cell line and MOI for downstream compound screening.

#### Step 4) Compound Library and Screening setup

Incorporation of appropriate controls, such as known antivirals, help benchmark assay performance. It is important to include sufficient uninfected wells and DMSO-treated infected wells. To minimize positional bias, avoid placing uninfected wells exclusively in the outermost rows or columns. We recommend using PLAID or other tools to effectively randomize and distribute controls and compounds across multiwell plates (Francisco Rodríguez et al., 2023). Consider including assay interference compounds, such as phospholipidosis inducers, as well as compounds of particular interest for your biological question. As a counter-screen, we propose to perform a DIPL assay to determine whether phospholipidosis is a confounding factor contributing to the observed antiviral activity.

#### Step 5) Analysis

Extracted morphological features can be analyzed in the provided Jupyter Notebooks to compute morphology scores and prioritize compounds that reverse virus-induced phenotypes. In addition, target enrichment analysis can be performed to gain mechanistic insights by linking phenotypic responses to known compound targets, aiding in the biological interpretation and prioritization of candidate hits.

Requirements to run the assay include: access to a biosafety laboratory appropriate for the viral pathogen’s containment level, high-throughput fluorescent microscope, adequate computational power and storage, a cloud-based platform with Jupyter Notebook, and CellProfiler installations for parallelized image analysis. We recommend referring to the original Cell Painting guidance provided in Bray et al., and Cimini et al., for detailed protocols on automated image acquisition, computational setup, and laboratory equipment requirements (Bray et al., 2016; Cimini et al., 2023).

## 4. Discussion

Host-targeting antiviral strategies offer a promising approach for addressing both current and future viral outbreaks, as they reduce the risk of viral resistance development, while providing broad-spectrum activity against multiple related pathogens. However, identifying compounds that effectively modulate host pathways to inhibit viral replication remains challenging using conventional screening approaches. In this study, we present a systematic drug repurposing strategy that addresses this challenge by combining morphological profiling with antibody-based detection of viral infectivity to identify host-targeting antiviral compounds. We conducted a comprehensive set of screens across 5,275 repurposing drugs, including a cell viability assay, an inhibition of cytopathic effects assay, a morphological profiling assay combined with antibody staining and a phospholipidosis counterscreen. We identified compounds that could reverse SARS-CoV-2–induced cellular phenotypes, targeting host pathways frequently hijacked by viruses, providing a robust pipeline for discovering host-directed therapeutics.

Our results demonstrate that SARS-CoV-2 infection induces characteristic and robust morphological changes in both Vero E6 and human A549^ACE2^ cells, with morphological features alone achieving near-perfect separation between infected and uninfected cells. We found that Cell Painting features capture virus-specific cellular effects, including strong endoplasmic reticulum (ER) changes and syncytia formation, consistent with the role of ER to support replication of SARS-CoV-2 (Williams et al., 2023), associated with the pathological effects observed in lung tissue (Buchrieser et al., 2020). While a few recent studies have begun to explore morphological profiling in the context of antiviral screening (Doijen et al., 2024; Mirabelli et al., 2021), our assay applies a broader, untargeted staining panel that captures seven major cellular compartments, which we believe provides a more detailed view on host cell responses and drug-induced effects.

To compare the hits from morphological profiling with those from conventional antiviral assays, we compared the results for three primary readouts: cytopathic effect (CPE), Cell Painting, and antibody-based detection. We found that Cell Painting and antibody-based assays showed substantial overlap in their hit sets, whereas the CPE assay yielded a distinct group of candidate compounds, likely due to its indirect measurement of antiviral activity, as well as the larger number and lower doses of compounds tested. Despite limited overlap at the compound level, all three assays resulted in hits covering a wide and largely overlapping range of mechanisms of actions. Notably, a few specific modes of action were enriched in individual assays, for example, NF-κB pathway inhibitors were only identified by the CPE assay, while Rho-associated kinase inhibitors were uniquely detected by Cell Painting, indicating that the assays carry complementary insights.

Among the antiviral hits identified from the three primary readouts, we observed a strong enrichment of compounds inducing drug-induced phospholipidosis (DIPL). This aligns with a report by Tummino et al., identifying DIPL as a frequent source of false positives in cell-based SARS-CoV-2 antiviral studies (Tummino et al., 2021). By incorporating a dedicated DIPL counter-screen, we were able to effectively deprioritize compounds whose antiviral activity could have been driven by phospholipid accumulation rather than true antiviral mechanisms. Phospholipidosis induction resulted in only subtle morphological changes that were not effectively captured by the aggregated morphology score. Consequently, this analysis approach failed to deprioritize phospholipidosis-inducing compounds. However, when analyzing subsets of non-infected cells, the morphological effects associated with phospholipidosis became more distinguishable, suggesting that this phenotype is detectable through Cell Painting images and could be identified, and potentially predicted, through more targeted analytical approaches.

Using pathway enrichment analysis of the annotated targets we found multiple host pathways frequently hijacked by viruses, including innate immune responses and kinases-related processes. Previous studies have shown that SARS-CoV-2 disrupts innate immune pathways for immune evasion (Gordon et al., 2020; Minkoff and tenOever, 2023). Moreover, kinase signaling is commonly modulated by viruses for cell cycle regulation, cytoskeleton reorganization, and immune evasion (Hajjo et al., 2023; Keating and Striker, 2012). Peptidyl-tyrosine modification, reflecting tyrosine phosphorylation by receptor tyrosine kinases (RTKs), was also highly enriched and has been shown important for viral replication of SARS-CoV-2 and influenza virus (Liang, n.d.; Sanchez et al., 2025). Additional enriched processes include oxidative stress responses, GPCR and MAPK signaling, each modulated during SARS-CoV-2 infection, with MAPK signaling hijacked by several viruses (Abdel Hameid et al., 2021; Gain et al., 2023; Gordon et al., 2020; Higgins et al., 2023; Keating and Striker, 2012). Together, these findings suggest that the antiviral activity of the compounds could arise from targeting host pathways critical to the viral life cycle, offering insight into important virus-host interactions and therapeutic opportunities.

The validated 74 compounds were prioritized based on assay performance, clinical phase, chemistry properties and route of administration to create a TPP directed towards rapid action during a pandemic (“Target product profiles,” n.d.). Among the top 15 ranked compounds, all but four have previously shown *in vitro* activity against SARS-CoV-2, with only two directly targeting viral proteins, indicating that host-modulating mechanisms may contribute to the antiviral activity of the remaining compounds (Kuzikov et al., 2021; Lin et al., 2018). Three compounds, balicatib, calpeptin and cathepsin inhibitor 1, target cathepsin L, a host protease involved in viral entry. Although cathepsin 1 inhibitor has not previously been reported active against SARS-CoV-2, its target cathepsin L, has clearly shown to facilitate SARS-CoV-2 entry via the spike protein. Zhao et al., demonstrated that inhibition of cathepsin L prevents infection both *in vitro* and *in vivo* (Zhao et al., 2021).

Among our top 15 hits, we identified both previously reported broad-spectrum antivirals and novel candidates. Serdemetan, BMS-566419, and metixene have not previously been reported active against SARS-CoV-2, suggesting three orally available, phase 2 compounds for further evaluation. Moreover, three of the 15 top compounds, sunitinib, emetine, and cycloheximide, are reported to exhibit broad-spectrum antiviral activity across multiple virus families. The kinase inhibitor sunitinib, is reported active against diverse viruses, including coronaviruses, (SARS-CoV-2, SARS-CoV, and MERS-CoV), hepatitis C virus, rabies virus, dengue virus, and ebola virus, with *in vivo* confirmation for some (Bekerman et al., 2017; Neveu et al., 2015; C. Wang et al., 2020; P.-G. Wang et al., 2020). Broad-spectrum activity is a common feature of host-directed antivirals and is often absent in direct-acting therapies. This reinforces the value of our screening strategy for identifying host-directed antivirals with a broad-spectrum applicability (Geraghty et al., 2021; Ji and Li, 2020).

Leveraging the single-cell resolution of morphological profiling, we demonstrate that the extracted features contain substantial mechanistic information. By analyzing specific subsets of the data, different biological questions can be answered, from target engagement to off-target effects. For example, phenotypic similarity to well-annotated compounds can be used to infer mechanisms of action for poorly annotated or novel molecules, or to identify undesired off-target effects (Dahlin et al., 2023).

One limitation of our approach is the reliance on cell lines that, while easy and permissive to the virus, may not fully capture the complexity of human airway biology. For example, Vero E6 cells are from non-human origin, and lack functional interferon signaling, which may compromise the relevance of host-targeting antiviral effects. Although we addressed this by validating hits in human A549^ACE2^ cells, future studies would benefit from using more physiologically relevant models, such as Primary Bronchial Epithelial Cells (Dinesh Kumar et al., 2022). Future implementations could also adapt the staining panel, for instance, replacing the antibody channel with an organelle-specific dyes to tailor the assay toward specific host-cell responses or toxicity endpoints.

In summary, we present a scalable and information-rich antiviral screening approach based on Cell Painting, capable of capturing antiviral effects, compound toxicity, and mechanistic insights within a single assay. Compared to conventional phenotypic methods such as CPE, it offers deeper biological resolution, broader applicability, and can be readily integrated with standard antibody-based protocols. We identified previously reported broad-spectrum antivirals, alongside novel candidates, whose targets are enriched in virus-host interaction pathways, supporting our host-targeting approach. We provide open access to an extensive collection of high-quality screening datasets, covering drug toxicity across multiple human cell lines, antiviral activity measured by cytopathic effect (CPE) assays, Cell Painting and antibody-based phenotypic readouts for 5,275 drugs in the repurposed library, complemented by DIPL assay on a subset of the library. This resource offers valuable biological data designed for broad reuse in antiviral research, phenotypic screening, and toxicology. We believe that this new approach along with the dataset, presents a valuable tool for future antiviral discovery and outbreak preparedness.

## Acknowledgements

This work was supported by the REMEDi4ALL consortium, which has received funding from the European Union’s Horizon Europe research and innovation programme under grant agreement No. 101057442. Views and opinions expressed are those of the authors only and do not necessarily reflect those of the European Union, who cannot be held responsible for them. This project additionally received funding within the SciLifeLab National COVID-19 Research Program and Knut och Alice Wallenbergs stiftelse (KAW 2020.0182; KAW 2020–0182, 2020-0032; KAW 2020.0241, V-2020-0699). O.S. acknowledges funding from the Swedish Research Council (Grants 2020-03731, 2020-01865, 2024-04576, and 2024-03566), FORMAS (Grant 2022-00940), Swedish Cancer Foundation (22 2412 Pj 03 H), and the Swedish strategic research programme eSSENCE. The authors acknowledge the support from the Chemical Biology Consortium Sweden (CBCS), Uppsala University and Karolinska Institutet; a national research infrastructure funded by the Swedish Research Council (dr.nr.2021-00179) and SciLifeLab.

The computations were enabled by resources provided by the National Academic Infrastructure for Supercomputing in Sweden (NAISS), partially funded by the Swedish Research Council through grant agreement no. 2022-06725, resources provided by Uppsala University at UPPMAX, and the Berzelius resource provided by the Knut and Alice Wallenberg Foundation at the National Supercomputer Centre.

Work with SARS-CoV-2 was performed at the BSL-3 Biomedicum Core Facility, Karolinska Institute. We thank Oscar de Capetillo’s research group for kindly providing us with the A549^ACE2^ cell line, as well as Charlotte Stadler and for donating cell lines. We thank Amelie Wenz for technical assistance with Cell Painting experiments and Martin Haraldsson and Annika Jenmalm Jensen for their chemical expertise.

## Author contributions

**Elin Asp**: Writing – original draft, investigation, methodology, validation, formal analysis, project administration. **Jonne Rietdijk**: Writing – original draft, investigation, methodology, validation, formal analysis. **Marianna Tampere**: Investigation, methodology, formal analysis, validation, project administration, supervision. **Hanna Axelsson**: Investigation, methodology, formal analysis, supervision. **Duncan Njenda**: Investigation, methodology. **Swapnil Potdar**: Formal analysis. **Adelinn Kalman**: Investigation, methodology. **Polina Georgieva**: Investigation, methodology, **Maris Lapins**: Methodology, formal analysis. **Flavio Ballante**: Data analysis. **Alicia Soler**: Critical data review, **Martin de Kort**: Critical data review, **Tero Aittokallio**: Data analysis, Critical data review. **Andrea Zaliani**: Critical data review. **Maria Kuzikov**: Critical data review. **Philip Gribbon**: Critical data review. **Donald Lo**: Critical data review. **Jordi Carreras-Puigvert**: Conceptualization, analysis and interpretations, supervision. **Brinton Seashore-Ludlow**: Conceptualization, writing - review & editing, supervision, methodology, formal analysis. **Ola Spjuth**: Conceptualization, supervision, analysis and interpretations, writing - review & editing. **Päivi Östling:** Conceptualization, supervision, writing - review & editing, funding acquisition.

## Appendix

### Supplementary methods

#### Cell culturing

Additional experiments with Vero cells and U-87 MG cells were maintained in Dulbecco′s Modified Eagle′s Medium (GIBCO), and MRC-5 cells were cultured in Minimum Essential Media. These were additionally supplemented with 10% FBS, 1% PS and maintained under standard conditions.

#### Virus infection

Zika virus (ZIKV, Strain H / PF / 2013) was propagated in C6/36 cells and viral titres determined in Vero cells. HCoV-229E (ATCC) were propagated and virus concentration determined in Huh-7 cells. Viral titers were determined by end-point dilution assay and immunofluorescence imaging of a virus-specific protein. Cells were infected with ZIKV at MOI 1 and incubated for 48h, and cells were infected with HCoV-229E at MOI 10 for 24h.

#### Antibody-based screen

The threshold for infection was defined as the mean plus three standard deviations from the virus NP-specific antibody intensity in the blank control. The set-off was determined on a plate-to-plate basis to avoid bias by batch effects and was checked for accuracy by checking for false positives in non-infected wells.

#### Drug-induced phospholipidosis screen

Wells with low cell count were excluded from the analysis and were defined as <100 cells/well at 24h, <200 cells/well at 48h, and <300 cells/well at 72h. Dose-response curves were generated for compounds with at least four doses and a minimum of DIPL at 50%. Compounds that did not meet this criterion were excluded from the curve fitting due to insufficient doses or DIPL response.

### Supplementary Files

Supplementary file 1: Viabiliy_Caco2

Supplementary file 2: Viability_Calu3

Supplementary file 3: Viability_HepG2

Supplementary file 4: Viability_U2OS

Supplementary file 5: Viability_VeroE6

Supplementary file 6: Primary Screen_CPE

Supplementary file 7: Primary Screen_Cell Painting_SARS-CoV-2 antibody readout

Supplementary file 8: ValidationScreen_Cell Painting_SARS-CoV-2 antibody readout

Supplementary file 9: % Drug-induced phospholipidosis

Supplementary file 10: Phospholipidosis_DoseResponse_24h

Supplementary file 11: Phospholipidosis_DoseResponse_48h

Supplementary file 12: Phospholipidosis_DoseResponse_72h

Supplementary file 13: Validated hits_hit prioritization

### Expanded View Figure legends

**Expanded View figure 1.**
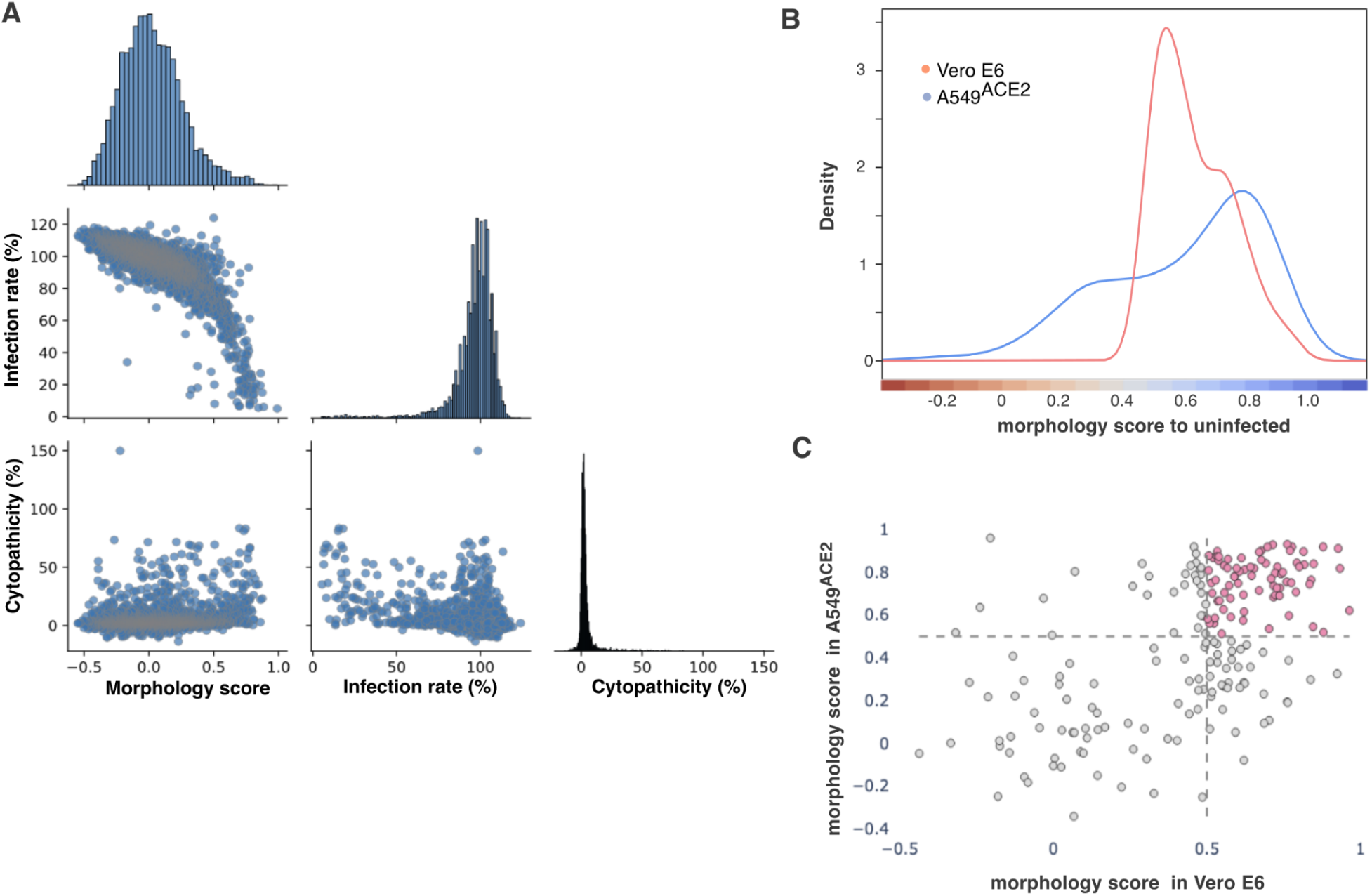
**A.** Pairwise plots of morphology score, inhibition of cytopathicity (%), and inhibition of viral infection for each perturbation (compound–dose combination). **B.** Density plots showing the distribution of morphology scores for the 150 prioritized compounds in Vero E6 and their respective scores in the A549^ACE2^ cell line. **C.** Scatterplot comparing morphology scores between Vero-E6 and A549^ACE2^ cells at matched doses for all 324 prioritized compounds. In pink are highlighted the compounds that achieved a morphology score of >0.5 in both cell models.

**Expanded View figure 2.**
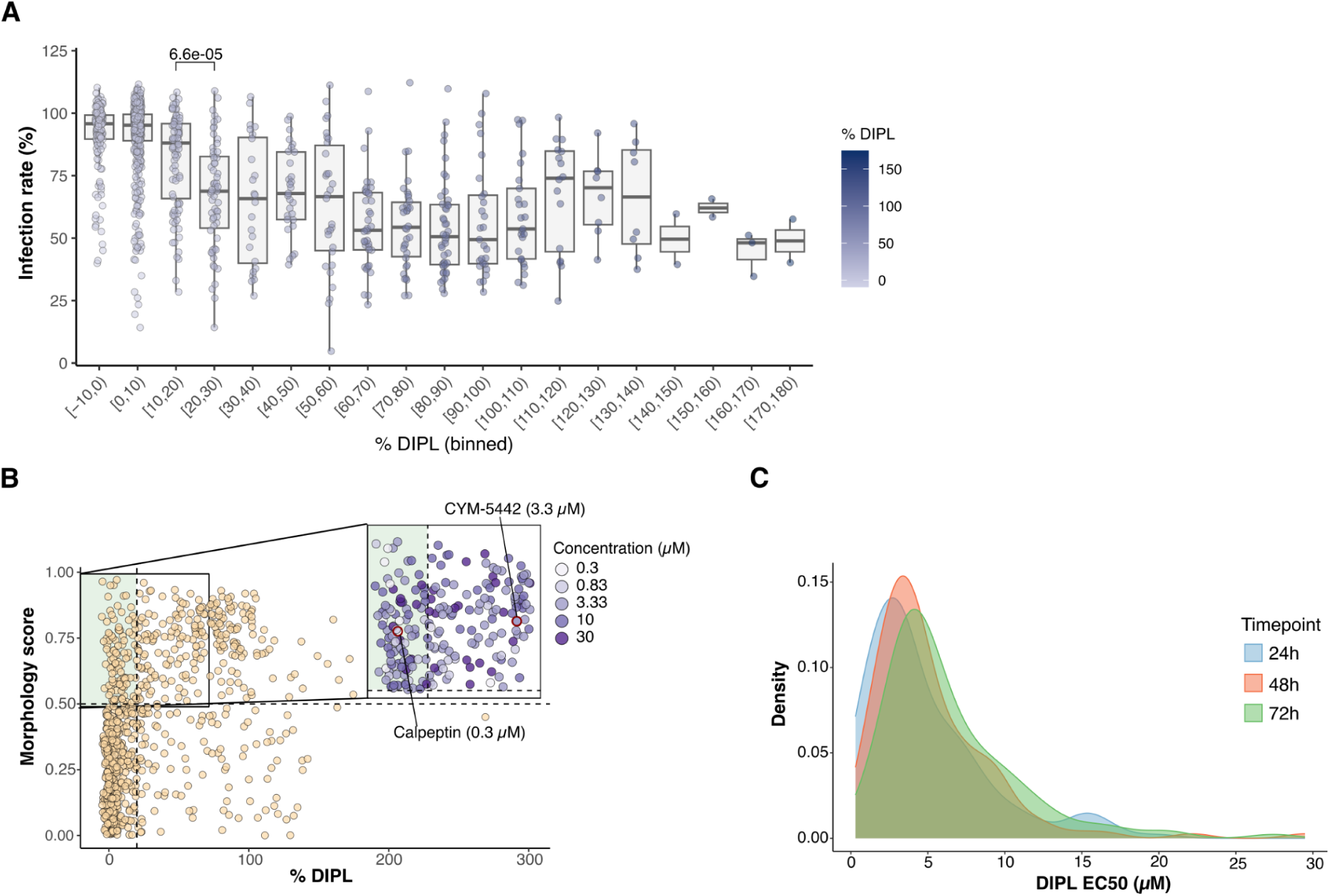
DIPL evaluation across timepoints and relationship with SARS-CoV-2 infection. **A.** SARS-CoV-2 infection rate across binned % DIPL values. Compounds were grouped into DIPL bins (10%-width), and infection rates are displayed as boxplots with individual data points overlaid. A significant decrease in infection rate was observed between 10-20% and 20-30% DIPL bins (Wilcoxon rank sum test, p = 6.6 × 10^−5^). **B.** A scatter plot showing the relationship between DIPL (x-axis, % normalized signal) and morphology score (y-axis) across tested concentrations of the 324 hit compounds. Each point represents a compound-concentration pair. Cut-offs show moderate DIPL level (<20%) and morphology score >0.5 for non-infected morphology. Compounds within the thresholds are indicated by a green-shaded area (n = 74). A close-up highlights compounds with non-infected morphology with varying DIPL levels, colored by concentration. Two compounds, CYM-5442 (3.3 μM) and Calpeptin (0.3 μM) are highlighted. **C.** Density plot showing EC_50_ values for DIPL across 24h, 48h, and 72h.

**Expanded View figure 3.**
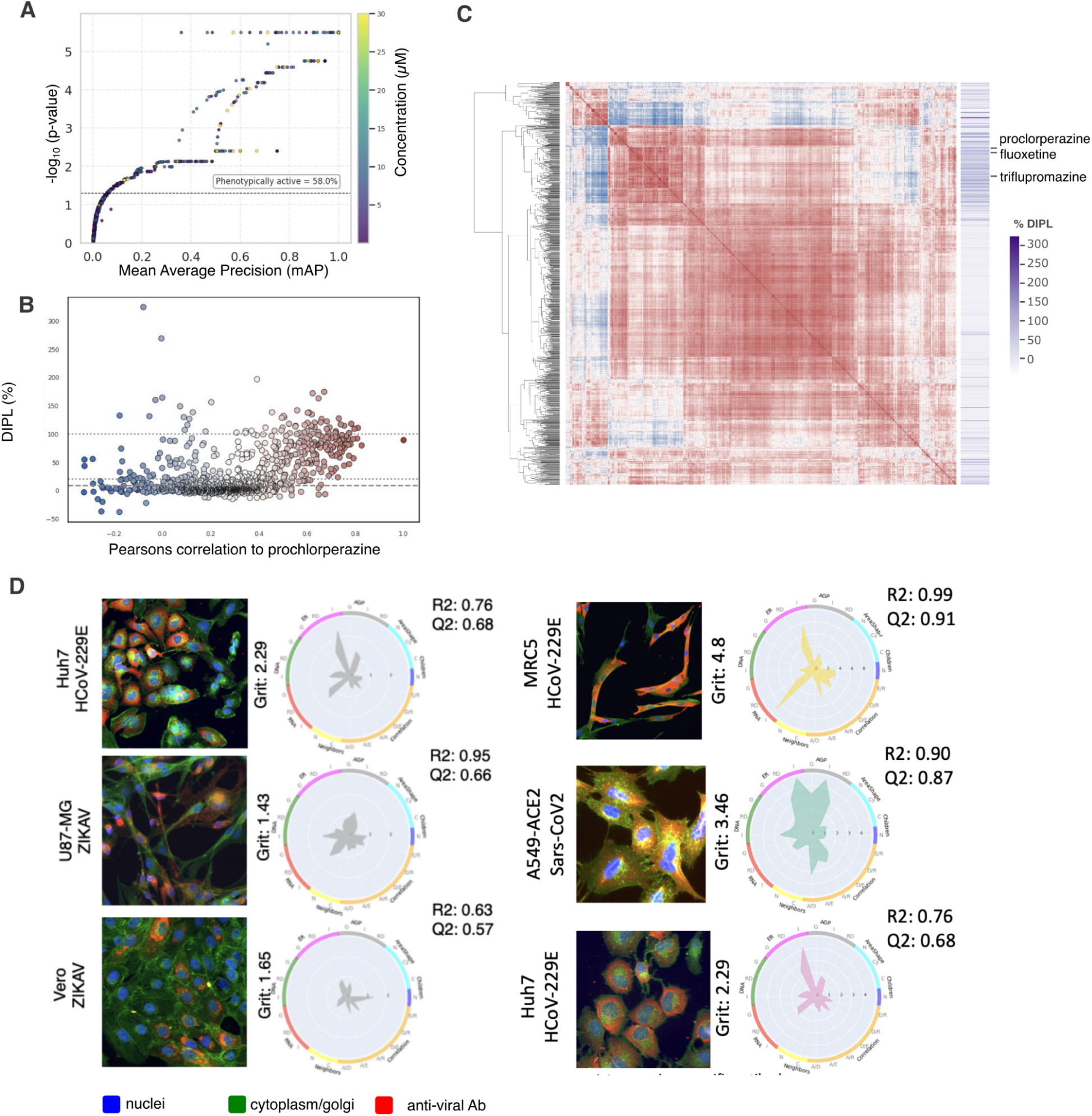
Morphological profiling of uninfected and infected cell populations. **A.** Mean average precision and –log₁₀(*p*-value) for uninfected morphological profiles. The dashed line indicates the significance threshold, the percentage of significantly altered morphological profiles is annotated above. **B.** Pearson correlation to fluoxetine hydrochloride versus percentage of phospholipidosis induction across compounds. **C.** Clustermap of Pearson correlations between compound-dose morphological profiles in uninfected cells. Highlighted compounds, prochlorperazine, triflupromazine, and fluoxetine, are shown. DIPL% values are represented as a continuous color bar. **D.** Representative Cell Painting images and radar plots for virus-infected cells in three cell line/virus combinations: MRC-5 with HCoV-229E, A549^ACE2^ with SARS-CoV-2 (used in this study), and Huh7 with HCoV-229E. Profile strength is summarized using R², Q², and grit scores. Fluorescence channels indicate nuclei (blue), virus-specific antibody (red), and cytoplasmic structures (green; including cytoskeleton, Golgi, and plasma membrane).

